# Allosteric activation of SARS-CoV-2 RdRp by remdesivir triphosphate and other phosphorylated nucleotides

**DOI:** 10.1101/2021.01.24.428004

**Authors:** Bing Wang, Vladimir Svetlov, Yuri I Wolf, Eugene V Koonin, Evgeny Nudler, Irina Artsimovitch

## Abstract

The catalytic subunit of SARS-CoV-2 RNA-dependent RNA polymerase (RdRp), Nsp12, has a unique NiRAN domain that transfers nucleoside monophosphates to the Nsp9 protein. The NiRAN and RdRp modules form a dynamic interface distant from their catalytic sites and both activities are essential for viral replication. We report that codon-optimized (for the pause-free translation) Nsp12 exists in inactive state in which NiRAN/RdRp interactions are broken, whereas translation by slow ribosomes and incubation with accessory Nsp7/8 subunits or NTPs partially rescue RdRp activity. Our data show that adenosine and remdesivir triphosphates promote synthesis of A-less RNAs, as does ppGpp, while amino acid substitutions at the NiRAN/RdRp interface augment activation, suggesting that ligand binding to the NiRAN catalytic site modulates RdRp activity. The existence of allosterically-linked nucleotidyl transferase sites that utilize the same substrates has important implications for understanding the mechanism of SARS-CoV-2 replication and design of its inhibitors.

**Highlights:**

- Codon-optimization of Nsp12 triggers misfolding and activity loss
- Slow translation, accessory Nsp7 and Nsp8 subunits, and NTPs rescue Nsp12
- Non-substrate nucleotides activate RNA chain synthesis, likely via NiRAN domain
- Crosstalk between two Nsp12 active sites that bind the same ligands

## INTRODUCTION

Respiratory RNA viruses pose an existential threat to humankind, the treat which proved to be the most refractory to modern sanitation and disease control measures that limited the spread of water-, food-, and blood-borne epidemics, such as cholera, plague, and AIDS (Piret and Boivin, 2020). All four pandemics of the XXI century, H1N1pdm09, SARS, MERS, and COVID-19 have been caused by respiratory RNA viruses, the latter three - by a β-coronavirus (CoV) that belongs to the vast order *Nidovirales* (Zhang et al., 2021). The ongoing COVID-19 pandemic led to a dramatic loss of human life and devastating economic and social disruptions around the world. The zoonotic origin of its causative agent, SARS-CoV-2, combined with the rapid rise of mutant strains within infected human population, and numerous instances of re-transmission to zoonotic hosts speak to its resilience as the persistent human pathogen, and the likelihood of the emergence of new β-CoV variants with pandemic potential (Case et al., 2021; Hu et al., 2021; Munir et al., 2020; Plante et al., 2021). Effective pandemic response measures, from the development of effective anti-virals to genomic surveillance, necessitate a detailed understanding of SARS-CoV-2 molecular and structural biology. Yet CoVs have been much less studied than other viral pathogens, in part owing to their extraordinary large genome size and complex biology (Gulyaeva and Gorbalenya, 2021).

Upon infecting human cells, CoV (+) strand RNA genome is translated to produce several non-structural proteins (Nsps) which are required for viral replication and gene expression (Snijder et al., 2016). Among these, SARS-CoV-2 Nsp12 plays a central role as a catalytic subunit of RNA-dependent RNA polymerase (RdRp). RdRps are broadly conserved among RNA viruses (Peersen, 2019) and are thus attractive targets for broad-spectrum antivirals (De Clercq and Li, 2016). Many nucleoside analogs identified as RNA synthesis inhibitors in other viruses have been actively pursued for retargeting against SARS-CoV-2.

However, CoV transcription machinery is quite unique: the transcribing RdRp associates with the replicative helicase Nsp13, proofreading exonuclease Nsp14/10, and several other viral proteins in a large membrane-bound replication-transcription complex (RTC) (Wolff et al., 2020); the RTC components are highly conserved among CoVs (Figure 1). Furthermore, unlike many well-studied single-subunit RdRps, a minimally active SARS-CoV-2 RdRp consists of Nsp12 and three accessory subunit, Nsp7 and two copies of Nsp8 (Chen et al., 2020b; Gao et al., 2020; Hillen et al., 2020) (7•8_2_•12; **Figure 2A**).

**Figure 1.**
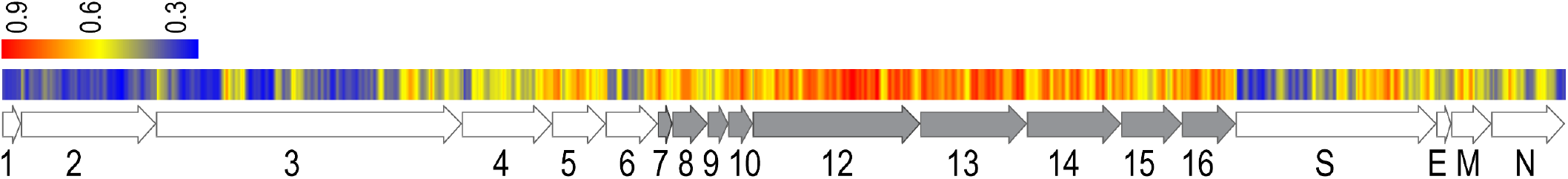
Conservation of amino acid residues in CoV genomes; only those proteins that are present in all *Coronaviridae* are shown; see **Dataset S1.** The Nsps are indicated by numbers; Nsp7-16 (shown in gray) that comprise the RTC are more conserved than structural (E, M, N, S) proteins and other Nsps.

**Figure 2.**
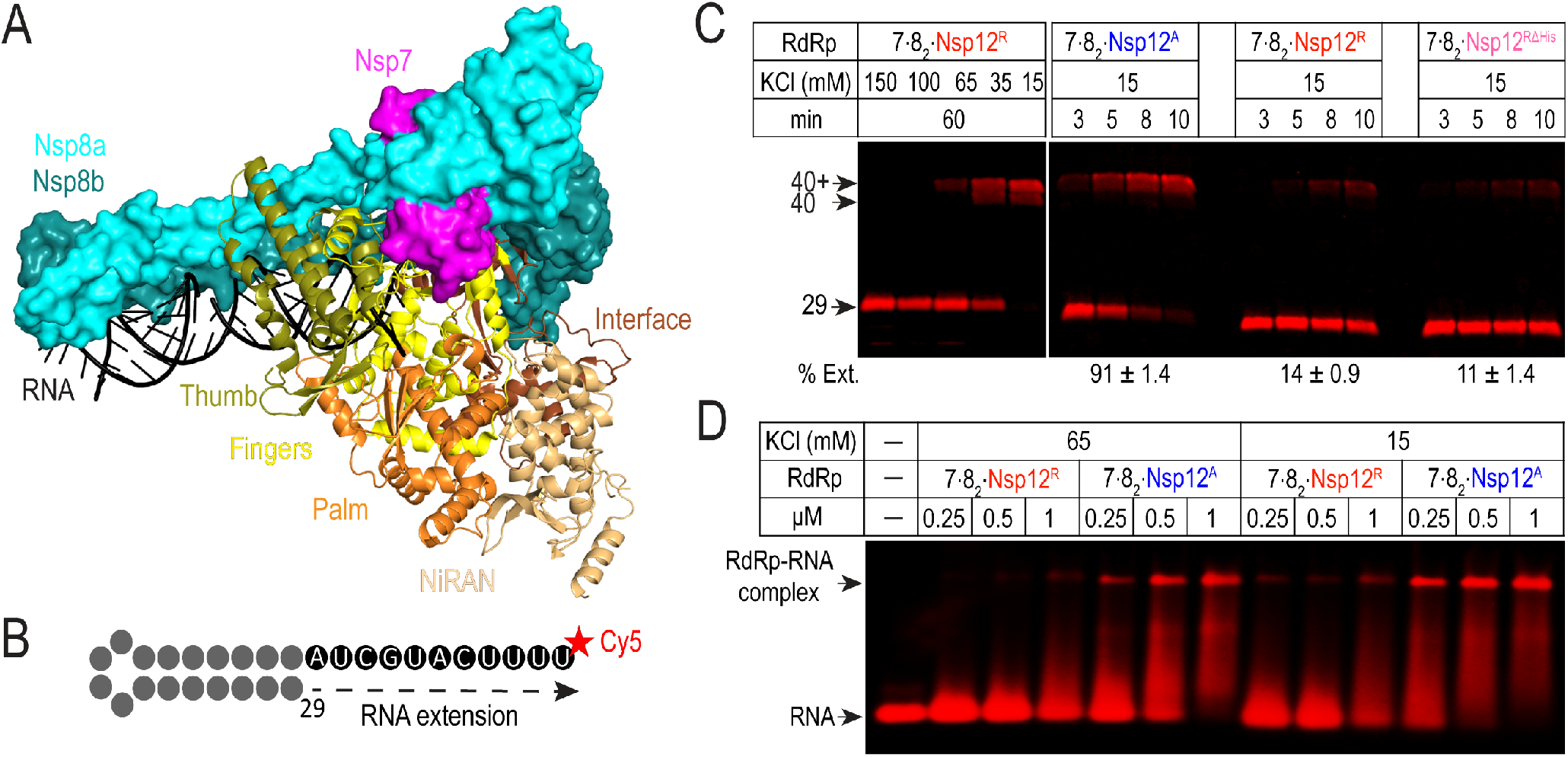
**A.** Transcribing RdRp. Nsp7 and Nsp8 are shown as surface, Nsp12 – as cartoon, with individual domains highlighted; PDB ID 6YYT. **B.** The 29-nt RNA hairpin scaffold is extended by RdRp to produce a 40-nt product; additional extension is thought to be mediated by Nsp8 after the completion of RNA synthesis (Tvarogova et al., 2019). **C.** RNA extension by RdRp at 37 °C under indicated conditions, permissive (15 mM KCl) conditions, removal of the His tag (ΔHis) does not increase Nsp12^R^ activity, but Nsp12^A^ expressed from an mRNA that retains rare codons is more active. Fractions of the extended RNA (% Ext.) at 10 min are shown (mean ± s.e.m.; n = 3). **D.** Interactions with the RNA hairpin scaffold analyzed by electrophoretic mobility shift assays. RdRps at indicated concentrations were incubated with 100 nM RNA at 37 °C for 5 minutes. Reactions were mixed with 10 × loading buffer (30 % glycerol, 0.2 % Orange G; Millipore Sigma) and run on a 3 % agarose gel in 1 × TBE on ice.

Nsp12 is a large (932 residues) multi-domain protein. In addition to the RdRp module composed of fingers, palm and thumb domains, Nsp12 contains a large <u>Ni</u>dovirus <u>R</u>dRp-<u>A</u>ssociated <u>N</u>ucleotidyl transferase (NiRAN) domain, which is connected to the fingers domain through an interface domain (**Figure 2A**). The NiRAN domain is endemic to nidoviruses and has been implicated in a range of activities, from RNA capping to protein-primed initiation of RNA synthesis (Lehmann et al., 2015). Recent report identified an accessory RNA-binding protein Nsp9 as the physiological target of NiRAN NMPylase and showed that this activity is critical for viral replication (Slanina et al., 2021); consistent with these findings, the NiRAN domain active site was observed to bind Nsp9 in a single-particle cryogenic electron microscopy (cryoEM) study (Yan et al., 2021), apparently in a catalytically-inactive arrangement. Thus, Nsp12 is a bi-functional enzyme with two active sites that transfer the NMP moiety to the 3’ end of the nascent RNA (AS1) or the N-terminus of Nsp9 (AS2). AS1 and AS2 utilize standard NTPs as substrates but may also accept a variety of ligands: AS1 readily incorporates remdesivir (Gordon et al., 2020) and favipiravir (Shannon et al., 2020b) monophosphates into RNA, and structural evidence suggests that AS2 may be similarly promiscuous (Chen et al., 2020b; Naydenova et al., 2021; Yan et al., 2021). Thus, the effects of NTPs and nucleoside analogs on both catalytic activities must be taken into account when interpreting experimental data and evaluating antiviral potential of lead molecules.

SARS-CoV-2 RdRp contains intrinsically disordered regions (IDRs) that undergo large context-dependent conformational changes, e.g., upon interaction with the product RNA (Hillen et al., 2020) or binding to ligands in the AS2 of NiRAN (Chen et al., 2020b; Naydenova et al., 2021; Yan et al., 2021). These inherent dynamic properties suggest that RdRp activity can be modulated, positively or negatively, by factors that control its folding. Here, we report that RdRp used in several structural and functional studies is poorly active because synonymous codon substitutions in Nsp12 designed to optimize its expression instead trigger its misfolding. We identify a region that contains a cluster of rare codons as important for the proper folding of Nsp12 and show that Nsp12 expression in a strain that encodes slow ribosomes and/or incubation with the accessory Nsp7/8 subunits increases RdRp activity. Finally, we show that nucleoside analogs that cannot be incorporated into RNA can nonetheless activate RNA chain extension, presumably through binding to AS2. Our findings have immediate implications for functional studies and identification of novel inhibitors of SARS-CoV-2 RdRp and highlight the need for improved mRNA recoding algorithms during rational design of other biotechnologically and medically important expression systems.

## RESULTS

### Nsp12 expressed from different coding sequences differ in activity and conformation

A rapidly growing collection of cryoEM structures of RdRp bound to different partners provides an excellent framework for understanding the mechanism of RNA synthesis and for the identification of novel RdRp inhibitors. As is the case with other systems, structural models require validation by functional studies that critically depend on the availability of robust expression systems and active RdRp preparations. Since structures obtained with RdRp produced in *Escherichia coli* (Chen et al., 2020b; Gao et al., 2020) and insect cells (Hillen et al., 2020) are similar, we used the *E. coli* expression platform (**Figure S1**) to initiate mechanistic studies of SARS-CoV-2 RdRp. For the sake of expediency, we used an Nsp12 expression vector described in (Gao et al., 2020); we refer to Nsp12 produced from this vector as Nsp12^R^ (where R indicates the laboratory where this plasmid was constructed) and Nsp7 and Nsp8-producing vectors constructed by us. Nsp12^R^ contains a non-cleavable C-terminal His_10_ tag, is relatively soluble when produced in *E. coli* and is easily purified under “native” (non-denaturing) conditions.

However, the 7•8_2_•12^R^ enzyme exhibited negligible activity on a number of different templates, including the optimal hairpin scaffold (**Figure 2B**) used by the Cramer group (Hillen et al., 2020), which could be extended only at very low salt. An extensive survey of purification schemes, RNA scaffolds, and reaction conditions failed to identify conditions that supported efficient primer extension, and the removal of the His tag, which was proposed to be detrimental (Dangerfield et al., 2020), did not increase activity under permissive (15 mM KCl) conditions (**Figure 2C**). In their follow-up study, Rao and colleagues reported similar results (Wang et al., 2020), prompting us to conclude that further improvement of activity of the 7•8_2_•12^R^ enzyme would be futile.

A robust enzyme activity is essential for functional studies - to rephrase the fourth commandment of enzymology (Kornberg, 2000), *thou shalt not waste clean thinking on dead enzymes*, and indispensable for the drug design and screening. Our survey of published reports failed to reveal an obvious reason for low activity - under similar reaction conditions, some RdRps were able to completely extend the RNA primer in minutes (Bera et al., 2021; Dangerfield et al., 2020; Hillen et al., 2020; Shannon et al., 2020b), whereas others failed to do so in an hour (Kirchdoerfer and Ward, 2019; Wang et al., 2020), regardless of the expression host. Idiosyncratic but reproducible variations in activity can arise from recombinant protein misfolding; indeed, co-expression of Nsp12 with cellular chaperons improves its activity (Dangerfield et al., 2020; Shannon et al., 2020a). A feasible source of this variability could lie in the coding mRNA itself: whereas all Nsp12s have the same amino acid sequence (ignoring the tags), their coding sequences (CDS) have been altered to match the codon usage of their respective hosts to maximize protein expression. This approach is routinely used by us and others (Gustafsson et al., 2004), yet protein function can be compromised even by a single synonymous codon substitution (Kimchi-Sarfaty et al., 2007; Liu, 2020), and protein expression in a BL21 RIL strain, which alleviates codon imbalance by supplying a subset of rare tRNAs and is thus commonly used to express heterologous proteins in *E. coli*, may hinder proper folding (Zhang et al., 2009). The loss of ribosome pausing at rare codons is thought to uncouple the nascent peptide synthesis from its folding, giving rise to aberrantly folded proteins (Liu, 2020; Liutkute et al., 2020; Pechmann and Frydman, 2013).

The viral *nsp12* mRNA is not optimized for translation in human cells (Finkel et al., 2020), whereas the *nsp12*^*R*^ codon usage matches that of highly expressed *E. coli* genes, suggesting that the *nsp12*^*R*^ CDS could be over-optimized. To evaluate this possibility, we designed an *nsp12*^*A*^ variant (where A indicates that it was constructed by IA) that contains more rare codons (**Figure S2**), including the regions that bear rare codons in the viral mRNA. We found that RdRp assembled with Nsp12^A^ had a much higher activity on the hairpin scaffold (**Figure 2C**). We also noted that Nsp12^A^ co-purified with nucleic acids; subsequent gel shift assays revealed that 7•8_2_•12^A^ readily bound the RNA hairpin, whereas 7•8_2_•12^R^ did not (**Figure 2D**).

To test whether Nsp12^R^ is misfolded, we used several approaches. *First*, we assessed the Nsp12 thermal stability using differential scanning fluorimetry (Abbott et al., 2017). We recorded melting temperatures (T_m_) of 41.3 °C for Nsp12^R^ and 47.3 °C for Nsp12^A^ (**Figure S3A**); T_m_ of 43.6 °C was reported for another *E. coli*-expressed Nsp12 (Peng et al., 2020). *Second*, we compared the intrinsic fluorescence spectra of Nsp12, which contains nine tryptophan residues expected to be sensitive to microenvironment (Chavez-Abiega et al., 2019). Nsp12^A^ and Nsp12^R^ exhibit similar emission peaks but the Nsp12^A^ intensity is two-fold higher (**Figure 3A**), suggesting that at least one Trp is more buried; the derivative spectra (**Figure S3B**) did not reveal any additional differences. These results show that Nsp12^A^ and Nsp12^R^ structures are different but cannot identify the altered regions.

**Figure 3.**
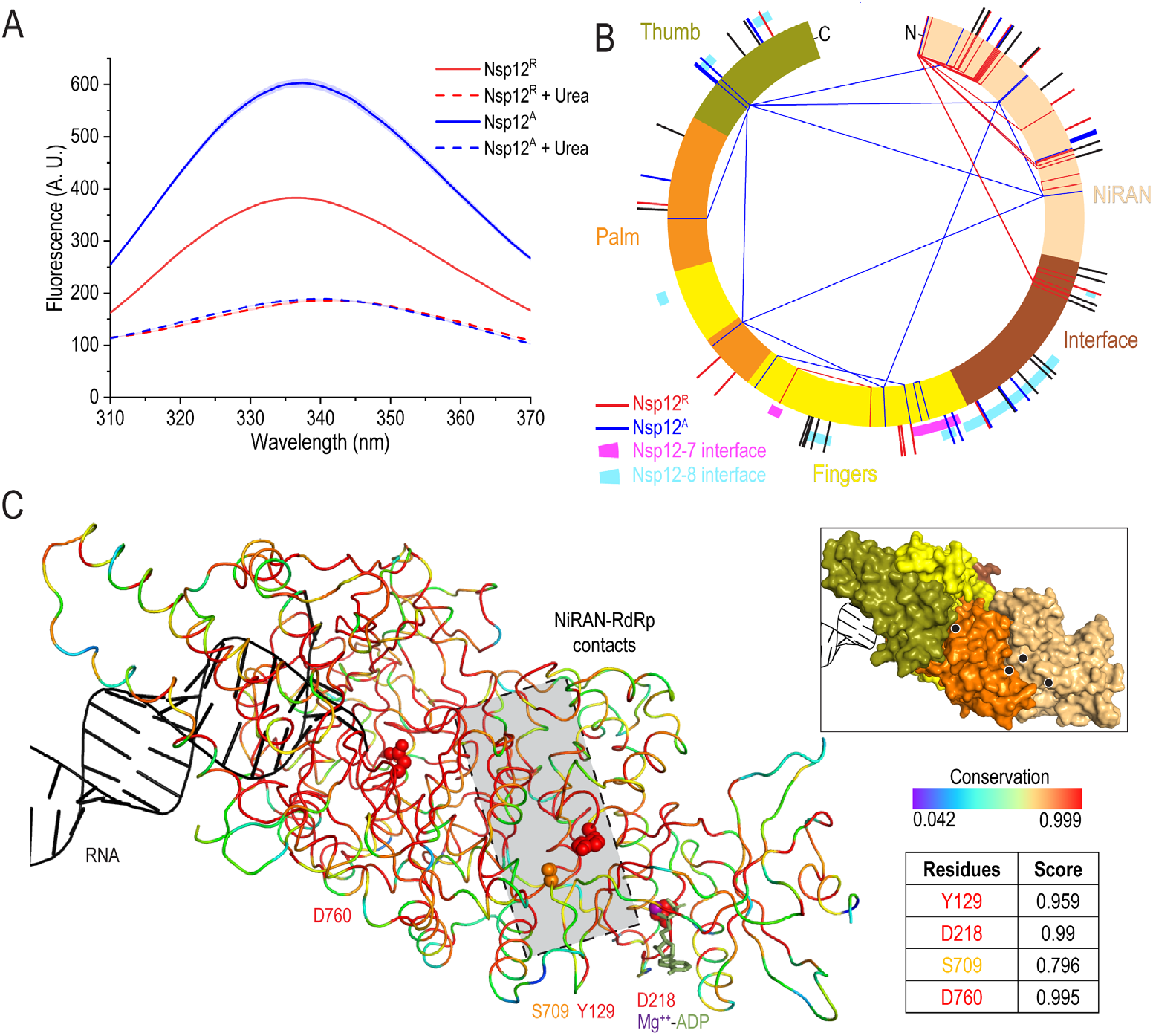
Differences between Nsp12^A^ and Nsp12^R^. **A.** Intrinsic tryptophan fluorescence of Nsp12 proteins; the spectra of denatured proteins confirm that their concentrations are identical. The mean and s.e.m. of triplicate measurements are shown as lines and shaded bands, respectively, in this and other figures. **B.** Mapping the EDC modifications. Lines show positions of monolinks (outside) and crosslinks (inside) mapped onto the Nsp12 schematic, with the domains colored as in panel A. Colors indicate differences in reactivity: residues in red were reactive only in Nsp12^R^; in blue – in Nsp12^A^; in black –in both proteins. Only high-confidence monolinks (<10^−5^) and crosslinks (<10^−3^) are shown; see **Dataset S2. C.** Conservation of the NiRAN/RdRp interaction surfaces mapped on the transcription complex structure; PDB ID: 6XEZ. Amino acid residues are colored according to their conservation. Key residues in AS1 (D760), AS2 (D218) and at the NiRAN/palm interface (Y129 and S709) are shown as spheres; ADP bound to AS2 is shown as sticks, the Mg^2+^ ion – as a purple sphere.

We next used a carboxyl and amine-reactive reagent EDC [1-ethyl-3-(3-dimethylaminopropyl)carbodiimide) to map solvent-accessible (surface) residues and intra-protein crosslinks by mass spectrometry. We observed significant differences in the accessibility of several regions centered at residues 150 (NiRAN domain), 415 (Fingers), 600 (Palm), and 850 (Thumb) and in crosslinking, particularly of the NiRAN domain (**Figure 3B**).

### Attenuated translation promotes active Nsp12 conformation

Although an overall abundance of underrepresented codons could slow translation, in many cases the ribosome must pause at one or more rare codons to promote folding of a “problematic” region (Yu et al., 2015; Zhou et al., 2015). There are more differences between the 2.8 kb mRNAs encoding Nsp12^A^ and Nsp12^R^ than there are similarities (**Figure S2**). The produced proteins also differ in their N- and C-termini (**Figure S1**) but, based on available structural data, an extra N-terminal glycine would not be expected to account for dramatic differences in EDC reactivity (**Figure 3B** and **Dataset S2**). Comparative analysis identified two regions that contained rare codon clusters in the viral and 12^A^ mRNAs, but not in 12^R^ (**Figure 3A** and **Figure S4**). We constructed chimeric proteins in which these 12^A^ segments were replaced with corresponding segments from 12^R^, generating proteins with identical amino acid sequences (**Figure 4A**). We found that while toggling of codons [143-346] between active and inactive Nsp12 variants did not alter the RdRp activity, a chimeric protein with codons [350-435] derived from the 12^R^ CDS was defective (**Figure 4B**). Together with the EDC modification patterns (**Figure 3D**), these data suggest that controlled translation of the 350-435 region is important for Nsp12 folding and that changes in contacts with Nsp7 (**Figure S5**), which are critical for RdRp activity (Hillen et al., 2020), could be partially responsible for the low activity of Nsp12^R^. Analysis of Nsp7/Nsp12 interactions by Trp fluorescence reveals that Nsp7 binds to both Nsp12 subunits and may favor a similar, Nsp12^A^-like state (**Figure S5**). In support of the “scaffolding” function of the accessory subunits (Peng et al., 2020), we found that preincubation of Nsp12^R^ with Nsp7 and Nsp8 led to an increased activity (**Figure 4C**).

**Figure 4.**
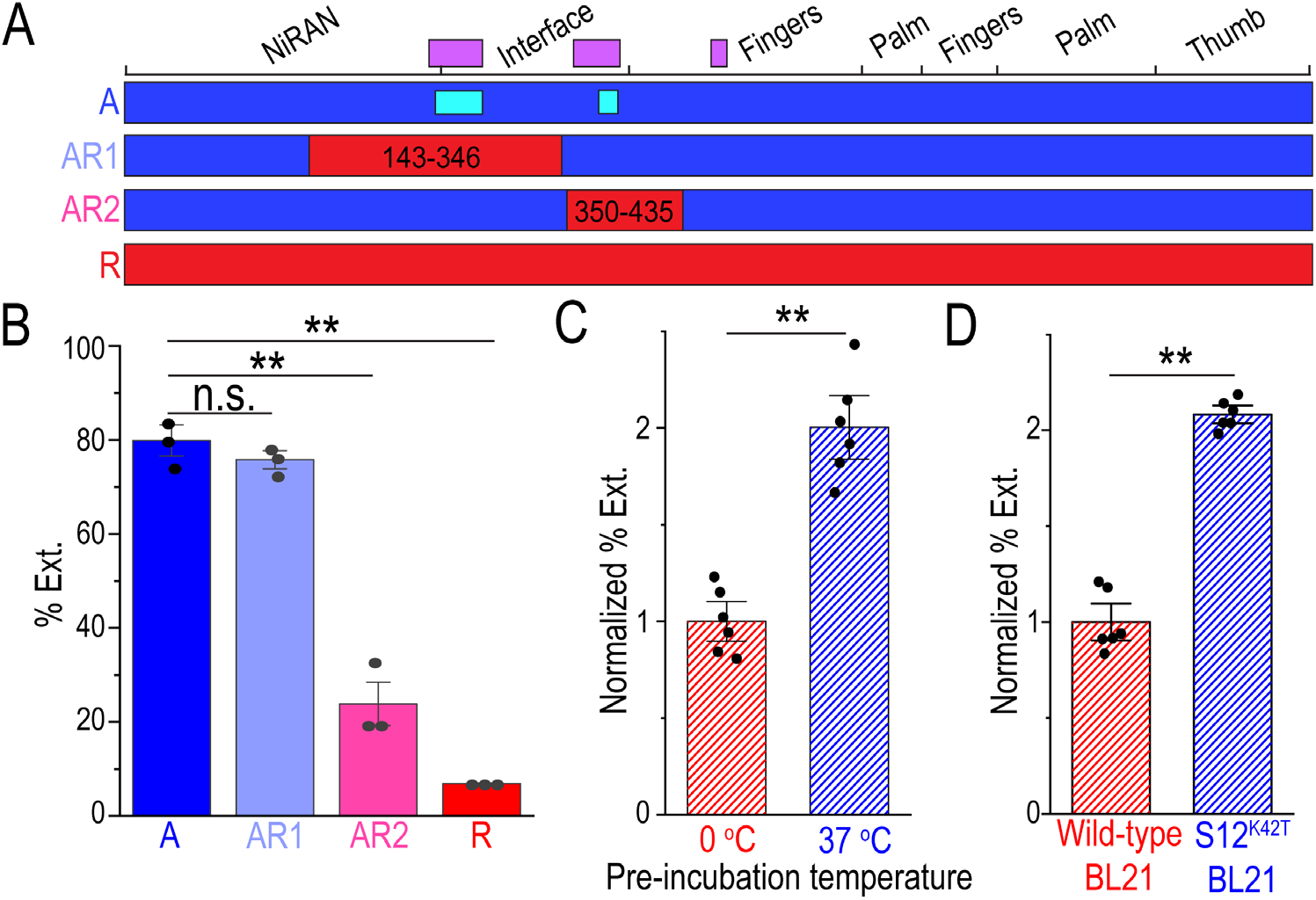
Determinants of the Nsp12 activity. **A.** Translational context around residue 400 is critical for the correct folding of Nsp12. SARS CoV-2 genomic *nsp12* RNA (with domain boundaries shown on top) contains clusters of rare codons (purple bars); only 12^A^ CDS has rare codons (cyan bars) at the corresponding positions. **B.** A chimeric Nsp^12-AR2^ protein is defective in RNA synthesis. **C.** Reactivation of Nsp12^R^ via 37 °C pre-incubation with the accessory Nsp7 and Nsp8 subunits to form the RdRp holoenzyme. (**D**) Translation by slow ribosomes yields a more active Nsp12. RNA extension is shown as mean ± S.E.M. and p value was calculated by unpaired two-tailed t-test. n.s., not significant; **, p < 0.01.

We next tested if slowing translation during protein expression would promote Nsp12 folding. We constructed a BL21 strain with a K42T mutant of the ribosomal protein S12, which shows ~ two-fold reduction in the translation rate (Ruusala et al., 1984), and compared the Nsp12^R^ protein purified from this “slow” BL21 variant to the protein purified from the wild-type BL21. We found that Nsp12^R^ purified from the mutant BL21 was ~ two-fold more active (**Figure 4D**), consistent with the favorable effect of attenuated translation.

### Allosteric RdRp activation by nucleotides

Our results show that Nsp12^A^ and Nsp12^R^ differ dramatically in the conformations/interactions of the NiRAN domain (**Figure 3B**). While NiRAN domain is not known to affect nucleotide addition, it interacts with the catalytic palm domain (Chen et al., 2020b; Hillen et al., 2020; Kirchdoerfer and Ward, 2019) and may modulate catalysis allosterically. The NiRAN domain is partially disordered in most unliganded structures of RdRp and transcription complexes, but becomes ordered upon binding of ADP-Mg^2+^, GDP-Mg^2+^ and PP_i_-Mg^2+^ to AS2 (**Figure 5A**) (Chen et al., 2020b; Naydenova et al., 2021; Yan et al., 2021). We hypothesized that, upon binding to nucleotides, the NiRAN domain may become less dynamic, favoring a more active RdRp conformation and leading to more efficient nucleotide addition. First, we compared RNA synthesis under “standard” conditions, in which RdRp is prebound to the RNA scaffold prior to the addition of NTP substrates, to the “NTP-primed” reaction in which the order of addition was reversed (**Figure 5B**). Our results show that preincubation with NTPs strongly potentiates Nsp12^R^ activity, an effect that could be mediated by the NiRAN domain.

**Figure 5.**
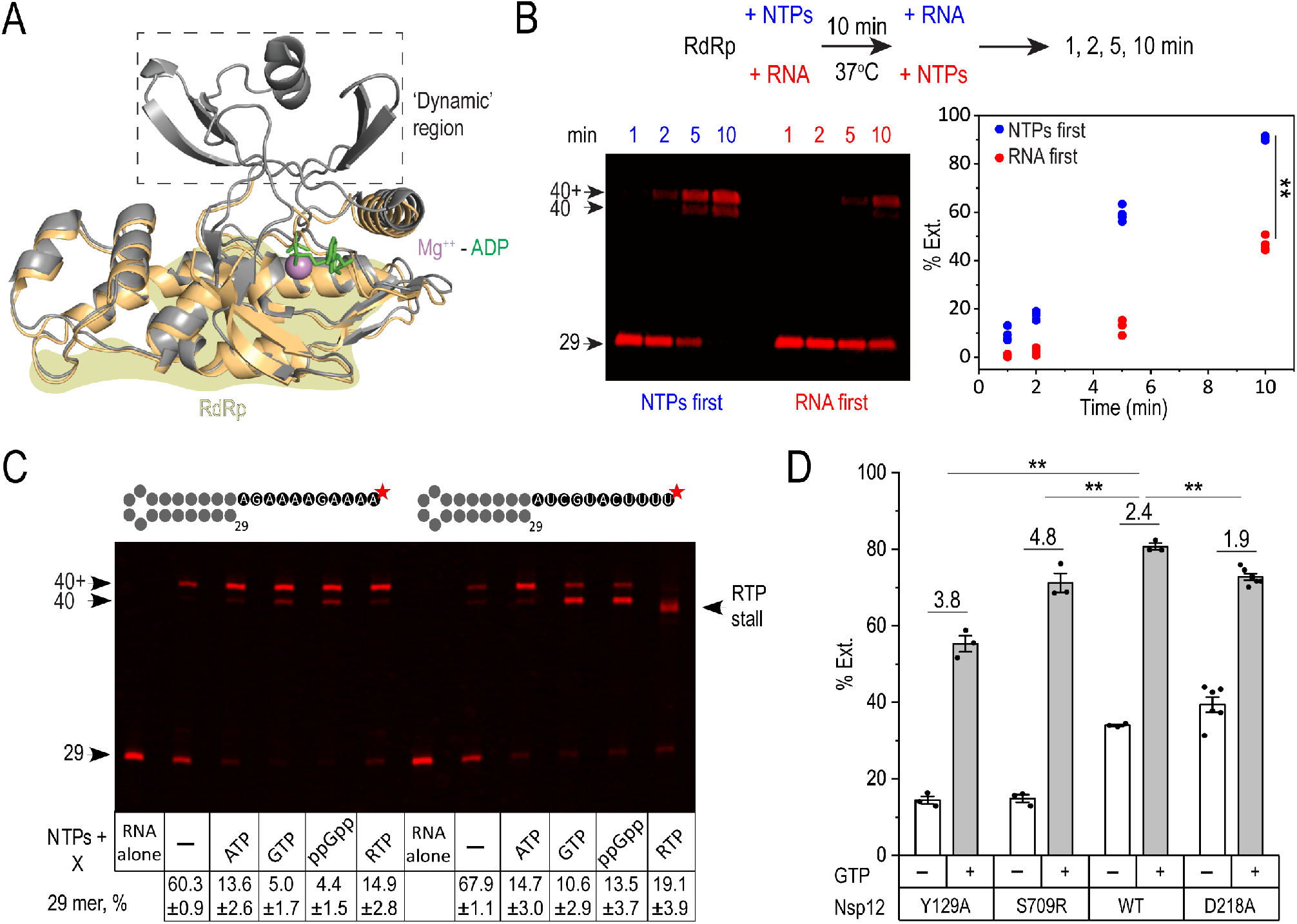
Allosteric activation of RdRp. **A.** An overlay of NiRAN domain structures in the absence (wheat) and in the presence (gray) of the bound ADP-Mg^2+^ (PBD IDs). The interface of the NiRAN domain and the RdRp domains is show. **B.** Activation of Nsp12^R^ holoenzyme by preincubation with NTPs. **C.** Activation of RNA synthesis by purine nucleotides (at 1 mM) on CU (left) and 4N (right) templates. **D.** Effects of Nsp12 substitutions on activation of RNA synthesis by 0.5 mM GTP; fold activation is shown above each set of bars. In B and D, RNA extension is shown as mean ± S.E.M. and p value was calculated by unpaired two-tailed t-test. n.s., not significant; **, p < 0.01.

Given that nucleotide binding to AS2 was found to remodel the NiRAN domain in the “active” Nsp12 (Chen et al., 2020b), we surmised that NTP-mediated activation should also occur in Nsp12^A^. To separate allosteric and direct effects of NTPs, we used a CU template which contains only purines in the transcribed region; as expected, the RNA was extended in the presence of CTP and UTP (**Figure 5C**); in addition to the run-off RNA (40 nt), a longer product that likely results from terminal transferase activity of Nsp8 was also observed; the latter activity prefers blunt over 3’ recessed ends and ATP as a substrate (Tvarogová et al., 2019). To assay hypothetical activation, we chose conditions under which less than 50 % of the scaffold was extended.

Consistent with the “allosteric” effects of non-templated nucleotides, transcription was activated in the presence of 1 mM ATP (> 4-fold) or GTP (> 10-fold) (**Figure 5C**). An apparent promiscuity of AS2 suggests that other nucleotides can substitute for ATP and GTP. To test this idea, we used a pause-promoting ATP analog remdesivir triphosphate (RTP) and an allosteric inhibitor of bacterial RNAP guanosine tetraphosphate (ppGpp). We found that the effects of RTP and ppGpp mimicked those of ATP and GTP, respectively, on the CU template. Activation was also observed on the 4N template, on which ATP, GTP and RTP but not ppGpp can be utilized as substrates; RMP incorporation led to RdRp stalling before reaching the end of the template, as expected (Gordon et al., 2020; Kokic et al., 2021). Consistent with the reported preference of the Nsp8 terminal transferase activity for ATP (Tvarogová et al., 2019), the fraction of the extended RNA is reduced in the presence of GTP and ppGpp (**Figure 5C**).

We hypothesized that RdRp-activating nucleotides likely act via binding to AS2 and stabilizing the RdRp/NiRAN interface. To evaluate this scenario, we substituted two conserved residues at the interface. Tyr129 in the NiRAN domain is nearly invariant among all CoV, whereas only small residues (Ser, Ala, Gly) are found at the 709 position in the palm domain (**Figure 3C**); we expected that these substitutions may compromise the interdomain contacts, making RNA synthesis more dependent on the state of the NiRAN domain. Consistently, we found that Y129A and Ser709R substitutions reduced RNA synthesis activity while potentiating activation by 0.5 mM GTP (**Figure 5D**).

The catalytic activity of the NiRAN domain was shown to be independent of the RdRp function: Nsp9 modification is fully proficient in an enzyme with substitutions in AS1 (Slanina et al., 2021). To determine if converse is true, we substituted Asp218, which binds to the AS2 catalytic Mg^2+^ ion (Chen et al., 2020b) and is critical for Nsp9 NMPylation and viral replication (Slanina et al., 2021), for Ala. This substitution did not compromise RNA synthesis, confirming that AS1 and AS2 are functionally independent, but modestly reduced GTP-dependent activation (**Figure 5D**) suggesting that, if the allosteric GTP binds to the NiRAN AS2, Asp218 does not significantly contribute to the nucleotide affinity. This observation is not entirely surprising because Asp residues are critical for the substrate positioning but make lesser contributions to the substrate binding in other viral polymerases (Gleghorn et al., 2011).

## DISCUSSION

Our results lead to two principal conclusions. *First*, SARS-CoV-2 Nsp12 depends on co-translational folding, facilitated by ribosome pausing, and on interactions with the accessory subunits to attain the active conformation. *Second*, the two nucleotidyl transfer catalytic sites in Nsp12, a unique property of nidoviruses, appear to be connected allosterically, with nucleoside analogs binding to the NiRAN AS2 activating RNA chain extension in the RdRp AS1.

### Pause-free translation inactivates RdRp

Our results demonstrate that over-optimized Nsp12^R^ mRNA produces a soluble but misfolded protein, in which RNA binding and catalytic activity (**Figure 2C, D**) are compromised. Notably, despite dramatic differences in their activities, all structures of SARS CoV-2 transcription complexes are similar (Chen et al., 2020b; Gao et al., 2020; Hillen et al., 2020), reflecting the bias introduced during cryoEM analysis, in which only a small fraction of “good” particles is picked for the analysis, e.g., 1% in a study of RdRp inhibition by remdesivir (Yin et al., 2020) . The preparation comprised of largely inactive enzymes is still amenable to the cryo-EM analysis, but would render many ensemble experiments futile. For example, a conclusion that SARS RdRp is more active than the SARS CoV-2 enzyme (Peng et al., 2020) is predicated on the assumption that both RdRps are properly folded. Even more critically, the inactive RdRp should not be used to screen potential inhibitors.

While recoding is routinely used to optimize heterologous protein expression (Gustafsson et al., 2004), the existence of rare codons in mRNAs encoding essential proteins indicates their important roles; for example, native non-optimal codons in IDRs are essential for function of circadian clock oscillators (Xu et al., 2013; Zhou et al., 2013). IDRs often serve as platforms for protein-protein interactions (Bugge et al., 2020), but can become trapped in unproductive states in the absence of their partners. Our analysis supports this scenario: an unstructured region that binds Nsp7 displays significant differential sensitivity to EDC (**Figure 2D**) and Nsp7 favors an “active-like” Nsp12 conformation (**Figure S5**). When added to the misfolded Nsp12, Nsp7/8 only modestly increase its activity (**Figure 3C**). However, as all Nsps are produced as a giant precursor in the virus-infected cells (Snijder et al., 2016), the accessory subunits could aid Nsp12 folding co-translationally, as apparently happens during their co-expression in *E. coli* (Dangerfield et al., 2020). Likewise, co-expression of *E. coli* RNA polymerase subunits suppresses assembly defects conferred by deletions in the catalytic subunits (Artsimovitch et al., 2003).

More broadly, our findings have implications for heterologous expression of countless other proteins. Although examples of deleterious synonymous substitutions have been reported, these examples were assumed to be outliers. In retrospect, optimization-induced misfolding may be far more prevalent than previously thought, with different recoding algorithms impacting the structure and activity of the resulting protein in vastly different ways. The importance of co-translational folding, particularly for large and dynamic proteins that contain essential mobile regions, emphasizes a need for integration of diverse approaches, from ribosome profiling to machine learning, during a rational design of coding sequences to avoid misfolding traps. Furthermore, this report provides much needed nuance in interpreting the results of sequence-based structural inferences, including homology modeling and *ab initio* predictions of protein structures and various AI approaches that utilize the implicit straightforward sequence-based structural determinism.

### Crosstalk between two catalytic sites of SARS-CoV-2 Nsp12

Decades of studies of viral RdRps focused on the mechanism of RNA synthesis and identification of nucleoside analogs that inhibit viral replication. During the Covid-19 pandemic, repurposing of the existing drugs targeting RdRp, justified by structural similarity among RdRp active sites (Peersen, 2019), became an urgent priority. Among these drugs, remdesivir received the most attention even though the estimates of its clinical effectiveness range from moderate to insignificant (Al-Abdouh et al., 2021; Kaka et al., 2021). RdRp readily uses RTP as a substrate in place of ATP and temporarily stalls downstream from the site of RMP incorporation (Gordon et al., 2020; Kokic et al., 2021); however, the proposed mechanisms of its action vary widely, from RdRp stalling to RNA chain termination to disassembly of the RdRp (Bravo et al., 2021; Olotu et al., 2021; Tchesnokov et al., 2020). It is presently unclear whether antiviral effects of remdesivir are due to delays in RNA synthesis or to errors in RNA messages, as is the case with another purine analog, favipiravir (Shannon et al., 2020b).

Although efforts aimed at identification of nucleoside analog inhibitors of RdRp are focused on AS1, it is clear that effects of nucleoside analogs on SARS-CoV-2 replication could be multifaceted. Nsp12 contains two active sites separated by more than 80 Å (**Figure 6**), both of which can bind NTPs and nucleoside analogs. Functions of AS1 and AS2 are largely independent: Nsp12 containing double substitutions in AS1 that abolish RNA chain extension is fully capable of Nsp9 NMPylation (Slanina et al., 2021), whereas D218A substitution in Nsp12 that abolishes NMPylation blocks viral replication (Slanina et al., 2021) but does not compromise RNA extension (**Figure 5D**). In addition, each subunit of the Nsp8 dimer can also bind NTPs (Tvarogová et al., 2019), and even though there is no structural evidence of nucleotide binding to Nsp8 and its terminal transferase activity could be post-transcriptional, SARS-CoV-2 RdRp has four nucleotide-binding sites.

**Figure 6.**
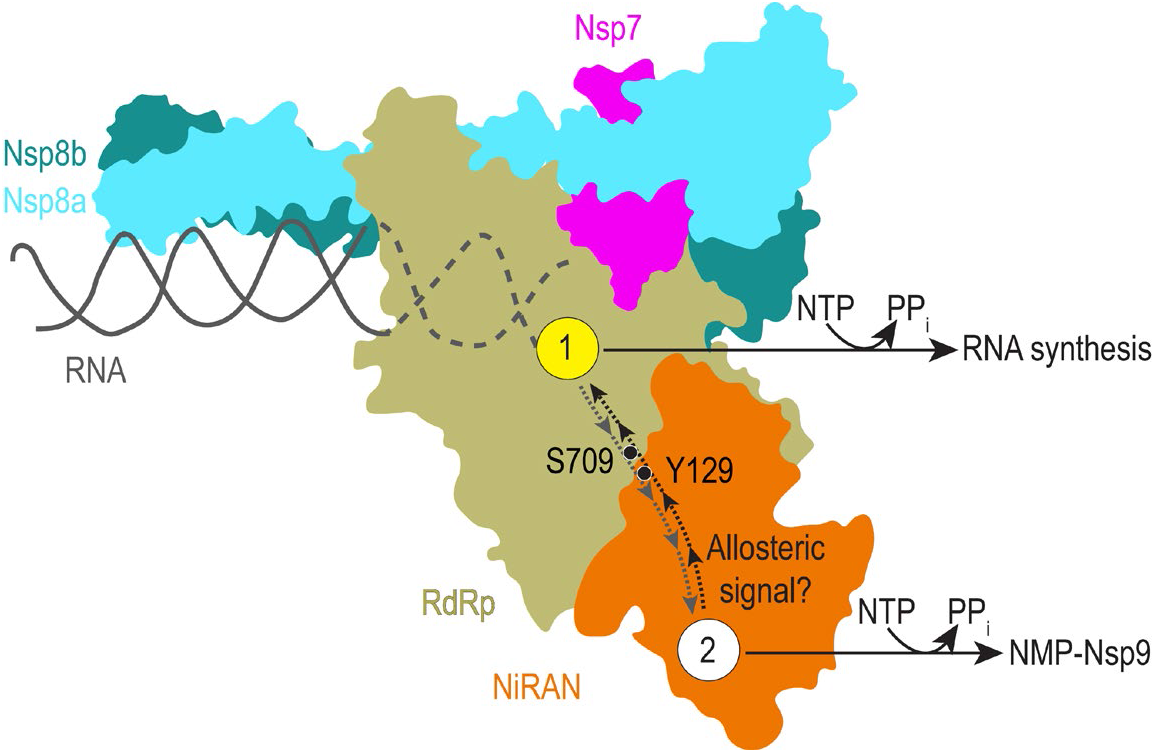
SARS-CoV-2 replication critically depends on two active sites in Nsp12 that mediate NMP transfer to RNA (AS1) and Nsp9 protein (AS2). Substrate (or inhibitor) binding to one site could be communicated to the other site through a highly conserved domain interface.

Thus, one cannot assume that the observed effect of a nucleotide is mediated via the “primary” AS1 – indeed, we show that RTP promotes RNA synthesis when it cannot be incorporated into RNA (**Figure 4C**), and ppGpp works even better. It is likely that competitive inhibitors binding in AS2 or noncognate ligands transferred to Nsp9 would inhibit replication. In the latter case, misincorporation could have more lasting effects because errors in the nascent RNA can be corrected by SARS-CoV-2 proofreading exonuclease Nsp14 (Shannon et al., 2020a).

We hypothesize that AS1 and AS2 are allosterically linked, enabling coordinated control. The NiRAN and palm domains form an extensive interface composed of highly conserved residues, including Y129 (**Figure 2C**). Upon binding to AS2, nucleotides induce NiRAN folding and lead to subtle changes at the domain interface (Chen et al., 2020b; Naydenova et al., 2021; Yan et al., 2021). We show that binding of nucleotides that cannot be incorporated into RNA potentiates RdRp activity (**Figure 5C**). We do not have direct evidence that this effect is triggered through their binding to AS2, but the effects of substitutions in the NiRAN domain (**Figure 5D**) and structural data (Chen et al., 2020b; Naydenova et al., 2021; Yan et al., 2021) support this model. While rigorous computational, structural, and biochemical analyses would be required to interrogate this hypothesis, it is already abundantly clear that, when considering the effects of various nucleotide analogs on viral RNA synthesis, their binding to AS2 (and perhaps AS3 and AS4) should not be ignored.

The open active sites of viral RdRp mediate binding of very diverse substrates, some of which have been developed into therapeutics (De Clercq and Li, 2016). Our findings that ppGpp activates RdRp similarly to GTP (**Figure 5C**) suggests that other nucleotide-binding sites in Nsp12 are also promiscuous. Furthermore, the interplay between binding of RTP to both catalytic and allosteric (relative to RNA synthesis) sites, and competition therein with cellular NTPs calls for a more nuanced interpretation of remdesivir mechanism, as does potential competition with ppGpp binding to allosteric (AS2) site. ppGpp biological activity has been long considered to be limited to bacteria and plastids, but its action as an alarmone has been recently demonstrated in human cells (Ito et al., 2020), raising possibility that ppGpp and other non-templating nucleotides may impact SARS-CoV-2 replication in host cells.

Allosteric control of SARS-CoV-2 RdRp invites interesting parallels with *E. coli* Qβ replicase, which also consists of four subunits, the phage-encoded RdRp (β-subunit) and three host RNA-binding proteins, translation elongation GTPases EF-Tu and EF-Ts and a ribosomal protein S1 (Takeshita et al., 2014). Similarly to Nsp7/8, EF-Tu and EF-Ts aid co-translational assembly of Qβ RdRp (Takeshita and Tomita, 2010); EF-Tu also forms a part of the single-stranded RNA exit channel, assisting the RNA strand separation during elongation, whereas S1 acts as an initiation factor (Takeshita et al., 2014). EF-Tu and EF-Ts binding to ppGpp modulates host translation (Rojas et al., 1984) and RNA synthesis by Qβ (Blumenthal, 1977), suggesting that RNA viruses from bacteria to humans may employ nucleotide analogs as sensors of cellular metabolism.

The SARS-CoV-2 RdRp subunit composition and dynamics resemble those of structurally unrelated bacterial RNA polymerases (RNAP). Bacterial enzymes are composed of 4-7 subunits and are elaborately controlled by regulatory nucleic acid signals and proteins that induce conformational changes in the transcription complex, as revealed by many recent cryoEM studies (Guo et al., 2018; Kang et al., 2018; Said et al., 2021). Notably, most natural and synthetic products that inhibit bacterial RNAPs alter protein interfaces or trap transient intermediates rather than block nucleotide addition and bind to many different sites (Chen et al., 2020a). As compared to simpler RdRps, SARS-CoV-2 enzyme, with several active sites and many conserved protein interfaces, could be an easier target for diverse small molecule that inhibit subunit or domain interactions or interrupt allosteric signals. Given the outsized importance of coronaviruses to human health, efforts to identify diverse inhibitors of RdRp, beyond RNA chain extension inhibitors, should be prioritized.

### Limitations of Study

The proposal that activating nucleotides (**Figure 5C**) exert their action upon binding to AS2 needs to be further investigated using systematic mutagenesis, and other phosphorylated nucleotide analogs should be evaluated for they ability to activate RNA extension. The proposed two-way crosstalk between AS1 and AS2 (**Figure 6**) needs to be investigated using *in vitro* assays of both enzymatic activities and nucleotides that have differential effects on both nucleotidyl transfer reactions.

## ACKNOWLEDGEMENTS

We are grateful to Georgi Belogurov, Venkat Gopalan, and Markus Wahl for suggestions, to Elena Kudryashova for help with data acquisition, and to Dmitri Svetlov for discussions and expert editing. This work was supported by a seed grant from The Ohio State University Office of Research to I.A.; the National Institutes of Health grants GM067153 (I.A.) and GM126891 (E.N.); and by Blavatnik Family Foundation and the Howard Hughes Medical Institute (E.N.). Y.I.W. and E.V.K. are supported by the Intramural Research Program of the National Institutes of Health (National Library of Medicine).

## AUTHOR CONTRIBUTIONS

Conceptualization, I.A.; Cloning, Protein Purification, and Biochemistry, B.W. and I.A.; Mass Spectrometry, V.S.; Bioinformatics analysis, Y.I.W; Funding Acquisition and Supervision, I.A, E.N., and E.V.K; Manuscript First Draft, B.W. and I.A. All authors contributed to finalizing the written manuscript.

## METHODS

### Construction of expression vectors

Plasmids used in this study are shown in **Figure S1**. The SARS-CoV-2 *nsp7/8/12*^*A*^ genes were codon-optimized for expression in *E. coli* and synthesized by GenScript and subcloned into standard pET-derived expression vectors under control of the T7 gene 10 promoter and *lac* repressor. The derivative plasmids were constructed by standard molecular biology approaches with restriction and modification enzymes from New England Biolabs, taking advantage of the existing or silent restriction sites engineered into the Nsp12 coding sequence. DNA oligonucleotides for vector construction and sequencing were obtained from Millipore Sigma. Sequence of all plasmids, including pET22a-Nsp12, were confirmed by Sanger sequencing at the Genomics Shared Resource Facility (the Ohio State University), and will be available upon request.

### Protein expression and purification

Nsp7/8 were overexpressed in *E. coli* XJB (DE3) cells (Zymo Research). Nsp12 variants were overexpressed in *E. coli* BL21 (DE3) cells (Novagen). Strains were grown in lysogeny broth (LB) with appropriate antibiotics: kanamycin (50 μg/mL), carbenicillin (100 μg/mL), chloramphenicol (25 μg/mL). All protein purification steps were carried out at 4 °C.

For Nsp7/8, cells were cultured at 37 °C to an OD_600_ of 0.6-0.8 and the temperature was lowered to 16°C. Expression was induced with 0.2 mM isopropyl-1-thio-β-D-galactopyranoside (IPTG; Goldbio) for 18 hours. Induced cells were harvested by centrifugation (6000 × *g*), resuspended in lysis buffer A (100 mM HEPES, pH 7.5, 300 mM NaCl, 5 % glycerol (v/v), 1 mM PMSF, 5 mM β-ME, 10 mM imidazole), and lysed by sonication. The lysate was cleared by centrifugation (10,000 × *g*). The soluble protein was purified by absorption to Ni^2+^-NTA resin (GE Healthcare), washed with Ni-buffer A (20 mM HEPES, pH 7.5, 300 mM NaCl, 5 % glycerol, 5 mM β-ME, 50 mM imidazole), and eluted with Ni-buffer B (20 mM HEPES, pH 7.5, 50 mM NaCl, 5 % glycerol, 5 mM β-ME, 300 mM imidazole). The eluted protein was further loaded onto a Resource Q ion-exchange column (GE Healthcare) in Q-buffer A (20 mM HEPES, pH 7.5, 5 % glycerol, 5 mM β-ME) and eluted with a gradient of Q-buffer B (20 mM HEPES, pH 7.5, 1 M NaCl, 5 % glycerol, 5 mM β-ME). The fusion protein was treated with TEV protease at 4 °C overnight, supplemented with 20 mM imidazole and was passed through Ni^2+^-NTA resin. The untagged, cleaved protein was loaded onto a Superdex 75 10/300 GL column (GE Healthcare) in Ni-buffer A. Peak fractions were assessed by SDS–PAGE and Coomassie staining. Purified protein was dialyzed into storage-buffer A (20 mM HEPES, pH 7.5, 150 mM NaCl, 45 % glycerol, 2.5 mM β-ME), aliquoted, and stored at −80°C.

For Nsp12, cells were cultured at 37 °C to an OD_600_ of 0.6-0.8 and the temperature was lowered to 16 °C. Expression was induced with 0.1 mM IPTG for 18 hours. Induced cells were harvested by centrifugation, resuspended in lysis-buffer B (100 mM HEPES, pH 7.5, 300 mM KCl, 5 % glycerol, 2 mM MgCl_2_, protease inhibitor cocktail (cOmplete, EDTA-free, Roche Diagnostics), 1 mM PMSF, 10 mM imidazole, 5 mM β-ME), and lysed by sonication. The cleared lysate was applied to Ni^2+^-NTA resin (GE Healthcare), washed with Ni-buffer C (20 mM HEPES, pH 7.5, 300 mM KCl, 5 % glycerol, 2 mM MgCl_2_, 5 mM β-ME, 0.1 mM PMSF) supplemented with 30 mM imidazole, and eluted with Ni-buffer D (20 mM HEPES, pH 7.5, 50 mM KCl, 5 % glycerol, 2 mM MgCl_2_, 5 mM β-ME, 0.1 mM PMSF, 300 mM imidazole). The eluted protein was further purified by Resource Q ion-exchange with Q-buffer C (20 mM HEPES, pH 7.5, 5 % glycerol, 2 mM MgCl_2_, 1 mM DTT) and Q-buffer D (20 mM HEPES, pH 7.5, 1 M KCl, 5 % glycerol, 2 mM MgCl_2_, 1 mM DTT). Then the fusion protein was treated with appropriate protease (TEV or SUMO protease) at 4 °C. After an overnight treatment, protein was supplemented with 20 mM imidazole and passed through Ni^2+^-NTA resin. The untagged protein was applied to the Superdex 200 increase 10/300 GL column (GE Healthcare) in SEC-buffer (20 mM HEPES, pH 7.5, 300 mM KCl, 5 % glycerol, 2 mM MgCl_2_, 1 mM DTT). Peak fractions were assessed by SDS–PAGE and Coomassie staining. Purified protein was dialyzed into storage-buffer B (20 mM HEPES, pH 7.5, 150 mM KCl, 45 % glycerol, 1 mM MgCl_2_, 1 mM DTT), aliquoted, and stored at −80°C.

### Expression by slow ribosomes

To test the effect of slow translation on Nsp12 activity, a derivative of BL21 containing a K42T substitution in the ribosomal protein S12 was constructed by P1 transduction from DEV3 *E. coli* strain (KL16 *lac5 strA2*; obtained from Kurt Fredrick) and selection on streptomycin (50 mg/L). This substitution reduces the translation rate ~ two-fold (Ruusala et al., 1984). Following sequencing of the *rpsL* gene to confirm the substitution, the slow BL21 was transformed with the plasmid encoding Nsp12^R^. The protein was purified as above.

### RNA extension assays

An RNA oligonucleotide (5’ -UUUUCAUGCUACGCGUAGUUUUCUACGCG- 3’) with Cyanine 5.5 at the 5’-end was obtained from Millipore Sigma (USA). Prior to the reaction, the RNA was annealed in 20 mM HEPES, pH 7.5, 50 mM KCl by heating to 75 °C and then gradually cooling to 4 °C. Reactions were carried out at 37 °C with 500 nM Nsp12 variant, 1 μM Nsp7, 1.5 μM Nsp8, 200 nM RNA, and 250 μM NTPs in the transcription buffer (20 mM HEPES, pH 7.5, 15 mM KCl, 5% glycerol, 2 mM MgCl_2_, 1 mM DTT). RNA extension reactions were stopped at the desired times by adding 2 x stop buffer (8 M Urea, 20 mM EDTA, 1X TBE, 0.2 % bromophenol blue). Samples were heated for 2 min at 95 °C and separated by electrophoresis in denaturing 9 % acrylamide (19:1) gels (7 M Urea, 0.5X TBE). The RNA products were visualized and quantified using Typhoon FLA9000 (GE Healthcare) and ImageQuant Software. RNA extension assays were carried out in triplicates. Means and standard error of the mean (s.e.m.) were calculated by OriginPro 2021 (OriginLab), and unpaired two-tailed t-test was performed using Excel (Microsoft).

### Electrophoretic mobility shift assays

RdRp (Nsp12: Nsp7: Nsp8 = 1: 2: 3; indicated concentrations in Figure 2D represent Nsp12 concentration) in 20 mM HEPES, pH 7.5, 65/15 mM KCl, 5 % glycerol, 2 mM MgCl_2_, 1 mM DTT were incubated with 100 nM 4N RNA at 37 °C for 5 minutes. Then reactions were mixed with 10 X loading buffer (30 % glycerol, 0.2 % Orange G; Millipore Sigma) and ran on a 3 % agarose gel in 1 X TBE on ice. The gel was visualized by Typhoon FLA9000.

### Activation of Nsp12^R^

To test the effect of holo RdRp formation, 5 μM Nsp12^R^ mixed with Nsp7/8 (10 μM/15 μM) in storage-buffer B was incubated at 0 °C or 37 °C for 15 min and then stored at −20 °C. RNA extension was performed as above, and the reaction was stopped at 8 min. To test the effect of NTPs, RdRp was first incubated with NTPs for 10 min at 37 °C, then 200 nM RNA was added to initiate the reaction; the final concentrations of RdRp (500 nM), RNA (250 nM) and NTPs (250 μM) were identical to those used in assays with the simultaneous addition of the RNA scaffold and substrates. The reaction was stopped by adding 2 x stop buffer at indicated times.

### Allosteric activation by nucleotides

A CU RNA hairpin (5’-AAAAGAAAAGACGCGUAGUUUUCUACGCG- 3’) labeled with Cyanine 5.5 at the 5’-end (Millipore Sigma) was annealed in 20 mM HEPES, pH 7.5, 50 mM KCl by heating to 75 °C and then gradually cooling to 4 °C. RdRp holoenzymes (500 nM wild-type or mutant Nsp12^A^, 1 μM Nsp7, 1.5 μM Nsp8; final concentrations) were mixed with ATP, GTP, RTP, or ppGpp at concentrations indicated in Figure 5 in 20 mM HEPES, pH 7.5, 15 mM KCl, 5% glycerol, 1 mM DTT and either 2 or 1 mM MgCl_2_ (with 1 or 0.5 mM activating nucleotide, respectively). Reactions were incubated for 5 min at 37 °C and RNA chain extension was initiated by the addition of 200 nM RNA and 100 μM CTP and UTP. Following 15 min incubation at 37 °C, reactions were stopped at the desired times by adding 2 x stop buffer (8 M Urea, 20 mM EDTA, 1X TBE, 0.2 % bromophenol blue).

### Tryptophan fluorescence

Tryptophan fluorescence spectroscopy was performed using F-7000 Fluorescence Spectrophotometer (Hitachi). The excitation wavelength was set at 280 nm and the emission spectra were recorded from 310 to 370 nm with a 5 nm slit width of excitation and emission. The scan speed was 240 nm/min. The temperature was maintained at 37 °C by a thermostatic water circulator (NESLAB RTE-7; Thermo Scientific). The samples were prepared in 20mM HEPES, pH 7.5, 65 mM KCl, 5 % glycerol, 2 mM MgCl_2_, 1 mM DTT. 1 μM Nsp12 and 2 μM Nsp7 was used to record the spectra of Nsp12 and Nsp7, respectively. To record the spectra of Nsp7•12, 1 μM Nsp12 was incubated with 2 μM Nsp7 at 37 °C for 15 min. To collect spectra of denatured proteins, 1 μM Nsp12 was incubated in 8 M urea at room temperature for 1 hour. Three independent measurements, each in three technical replicates, were performed. The same results were obtained with proteins purified three months apart. Means, s.e.m. and second derivatives of the emission spectra were calculated by OriginPro 2021.

### Conservation analysis

To assess relative conservation of coronavirus proteins, 3,309 diverse coronavirus genomes, representing alpha-, beta-, gamma-, and deltacoronavirus genera were downloaded from GenBank in May 2020. High-quality CDS (containing no more than 32 contiguous codons with ambiguously bases) were translated into five (poly)proteins, conserved across all coronaviruses: orf1ab, S, E, M, and N. Alignments of five ORFs were produced using the MUSCLE program (Edgar, 2004). For each alignment column homogeneity and the weighted fraction of non-gap characters (both ranging from 0 to 1) were calculated as described previously (Esterman et al., 2021). The product of these two values was used as the conservation index (ranging from 0 to 1). For the whole-genome pan-*Coronaviridae* conservation map, only consensus positions (those with the fraction of gaps below 0.5) were used.

### EDC modification and mass spectrometry

~0.5 mg/mL of Nsp12 in 20 mM HEPES, pH 7.5, 50 mM KCl, 2 mM MgCl_2_, 1 mM DTT was mixed with fresh prepared EDC (*N*-(3-Dimethylaminopropyl)-*N’*-ethylcarbodiimide hydrochloride; Sigma). EDC was added to a final concentration of 2 mM and the reaction was performed at room temperature for 30 min. The reaction was quenched with 50X molar excess Tris-HCl (pH 7.5) for 5 min. Cross-linked protein samples were separated using SDS-PAGE, the protein bands were stained with GelCode Blue, and tryptic peptides were generated using In-Gel Tryptic Digestion Kit; peptides were purified using Pierce 10 μL C18 tips (all Thermo Fisher). Peptides were analyzed in the Orbitrap Fusion Lumos mass spectrometer (Thermo Scientific) coupled to an EASY-nLC (Thermo Scientific) liquid chromatography system, with a 2 μm, 500 mm EASY-Spray column. The peptides were eluted over a 180-min linear gradient from 96% buffer A (water) to 40% buffer B (ACN), then continued to 98% buffer B over 20 min with a flow rate of 200 nL/min. Each full MS scan (R = 60,000) was followed by 20 data-dependent MS2 (R = 15,000) with high-energy collisional dissociation (HCD) and an isolation window of 2.0 m/z. Normalized collision energy was set to 35. Precursors of charge state 2-6 and 4-6 were collected for MS2 scans; monoisotopic precursor selection was enabled and a dynamic exclusion window was set to 30.0 s. The resulting raw files were searched in enumerative mode with pFind3(Chi et al., 2018) in open search mode against Nsp12 sequence, the inferred modifications over 1 % cutoff were used as “variable” modifications in the subsequent pLink2 search. The same files were then searched in cross-link discovery mode using pLink2(Chen et al., 2019) against Nsp12 sequence, using [EDC] as the cross-linking reagent, trypsin as the enzyme generating the peptides, and variable modifications set as inferred by pFind3.

## SUPPLEMENTARY FIGURES

**Figure S1.**
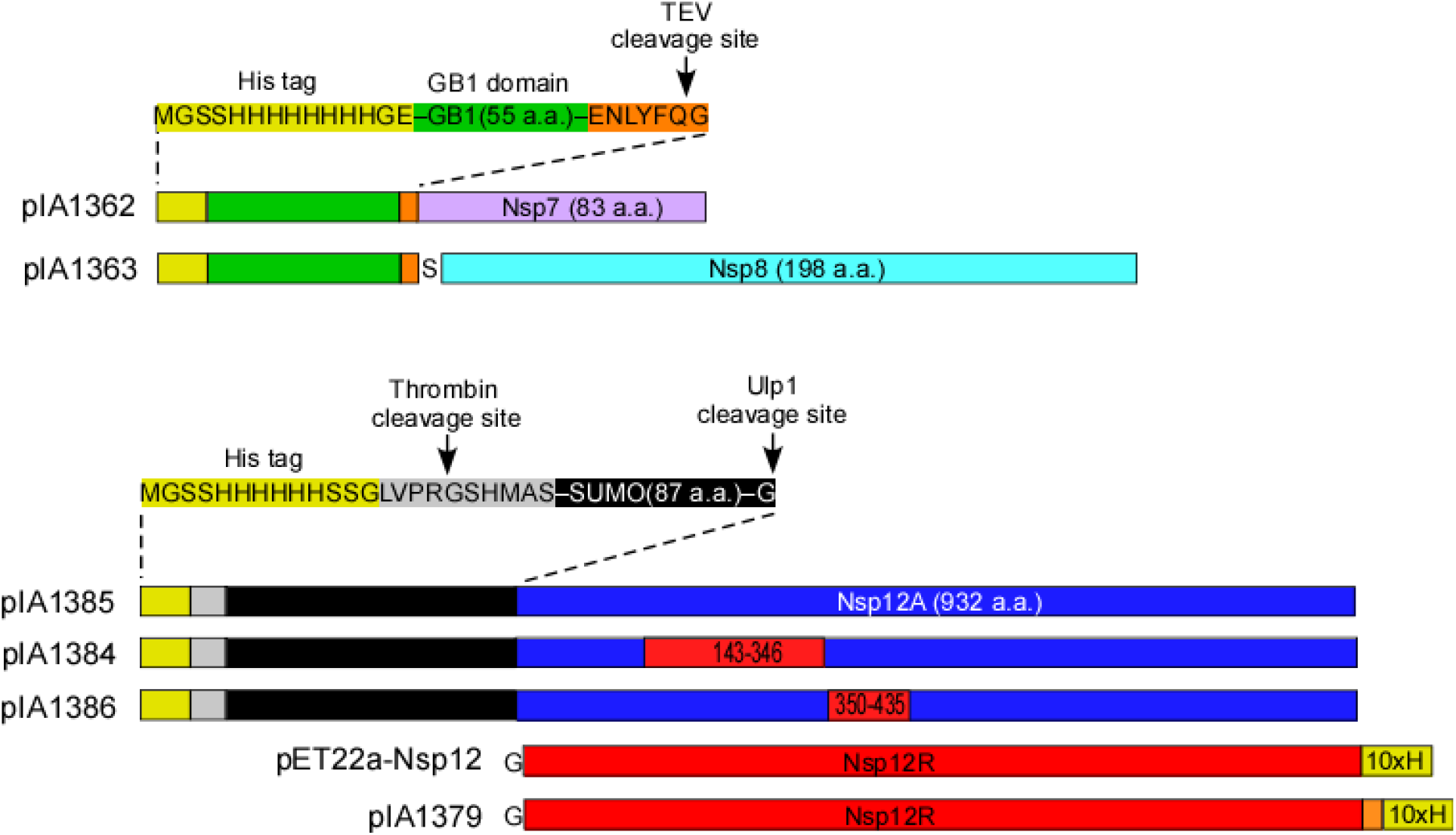
Expression constructs used in this study. pET22a-Nsp12 plasmid has been described by Dr. Rao and colleagues in Gao *et al.*(Gao et al., 2020); we refer to this variant as Nsp12^R^. In our group, genes encoding SARS-CoV-2 Nsp7, Nsp8, and Nsp12 proteins were obtained from GenScript; standard GenScript codon-optimization algorithm was used and additional silent restriction sites were designed to facilitate subsequent mutagenesis; this variant is referred to as Nsp12^A^. In addition, solubility and purification tags were added, together with the proteolytic cleavage sites. The synthetic cassettes were cloned into pET-based expression vectors under the control of the T7 gene 10 promoter and *lac* repressor. The relevant features of the expression vectors are shown. In vectors with N-terminally His-tagged Nsp12 variants (12^A^ and chimeras), the Ulp1-mediated cleavage generates the “authentic” N terminus of the protein, a feature that could be critical for RdRp activity(Gohara et al., 1999). Our data indicate, however, that additional sequences at N- and C-termini do not impair Nsp12 activity; we found that Nsp12 with the N-terminal His-SUMO tag is active, although we have not rigorously compared the activities of RdRps assembled with Nsp12 before and after cleavage by Ulp1, and that the removal of the C-terminal His tag does not increase the activity (**Figure 2C**); we note that the TEV-generated C-terminus in this and other cases has additional residues comprising the TEV recognition sequence. We also found that Nsp12^T^, expressed from a plasmid similar to pIA1385 that was constructed in the Tuschl lab for studies of the RdRp-Nsp13 helicase complexes(Chen et al., 2020b) (obtained from Addgene, #159107), was as active Nsp12^A^ under our assay conditions (not shown). To produce Nsp12^T^, the viral *nsp12* RNA was reverse transcribed and expressed in the BL21 RIL strain, which contains extra copies of the *argU*, *ileY*, and *leuW* rare tRNA genes. While comparable codon frequency measurements are not available for the RIL strain, it does not carry all rare tRNAs required for the efficient translation of the viral *nsp12* mRNA, suggesting that Nsp12^T^ codon usage is suboptimal. Since this plasmid does not allow for the expedient construction of mutationally-altered Nsp12 variants, we used pIA1385 instead.

**Figure S2.**
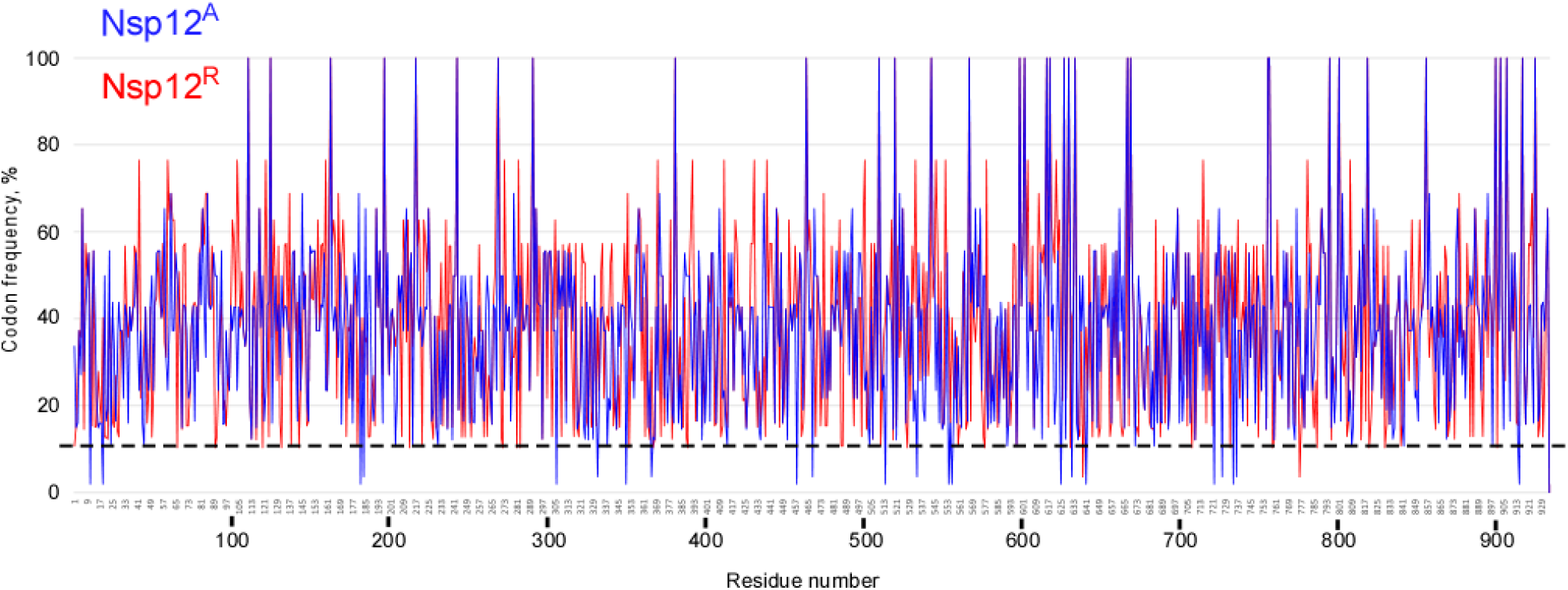
Differences in codon frequencies between Nsp12^A^ (blue) and Nsp12^R^ (red) CDSs. A dashed line indicates a 10 % frequency cut-off. In Nsp12^R^ mRNA, 2 codons fall below this threshold, as compared to 22 in Nsp12^A^ mRNA.

**Figure S3.**
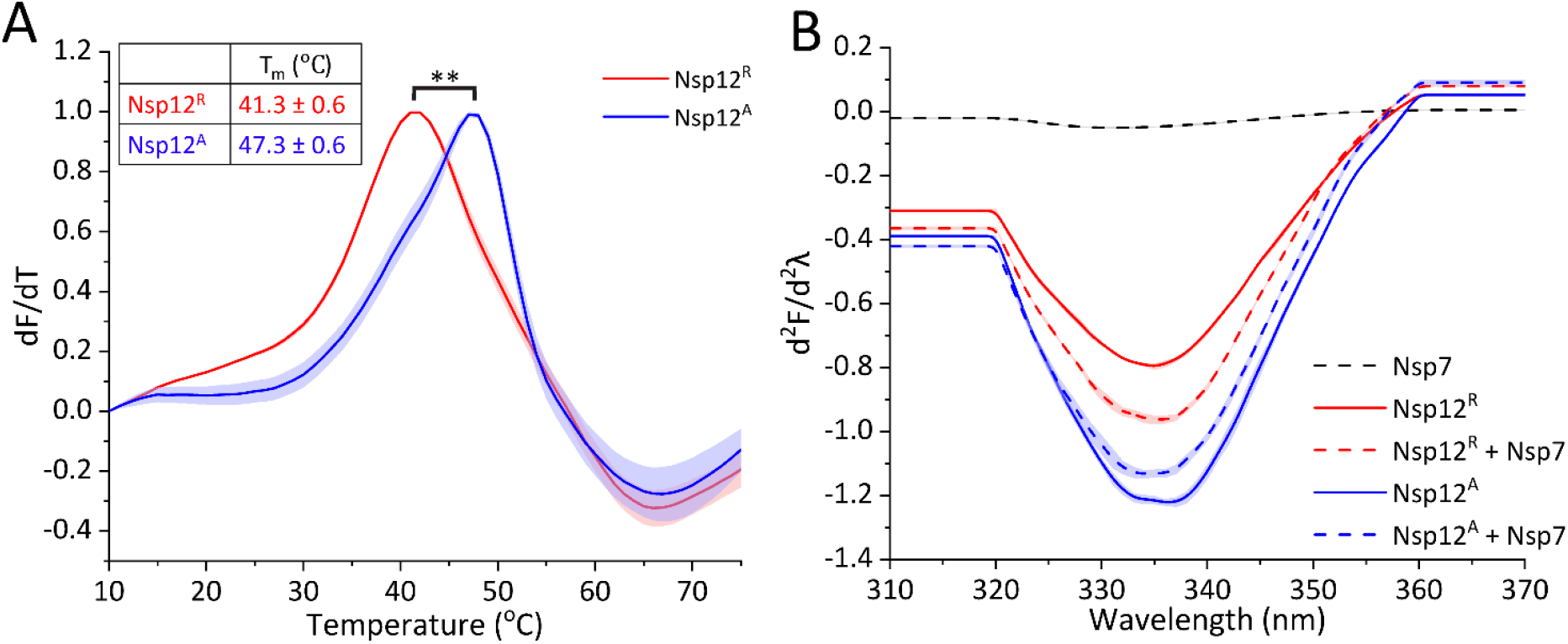
Structural differences between Nsp12^A^ and Nsp12^R^ proteins. **A.** Differential scanning fluorimetry (DSF) was conducted as reported previously(Schwebach et al., 2020). Proteins were diluted to 3 μM in 20 mM HEPES, pH 7.5, 150 mM KCl, 40 % glycerol, 1 mM DTT, 2 mM MgCl_2_ in the presence of 1:5,000 dilution of Sypro Orange dye (Invitrogen). Change in fluorescence of the dye, which preferentially binds to protein hydrophobic regions exposed upon heat-induced unfolding, was measured in triplicates at a rate of 2 °C/min using a CFX Real-Time PCR Detection System (Bio-Rad). The data are plotted as the first derivatives of the fluorescence signal versus temperature; three individual spectra are shown for each Nsp12 variant. The melting temperatures (T_m_) were determined as the maximum of the first derivative of each normalized experimental curve and expressed as the mean ± s.e.m., and the p value was calculated by unpaired two-tailed t-test. **, p < 0.01. **B.** Second derivatives of Trp emission spectra shown in **Figure 3A and S5B**.

**Figure S4.**
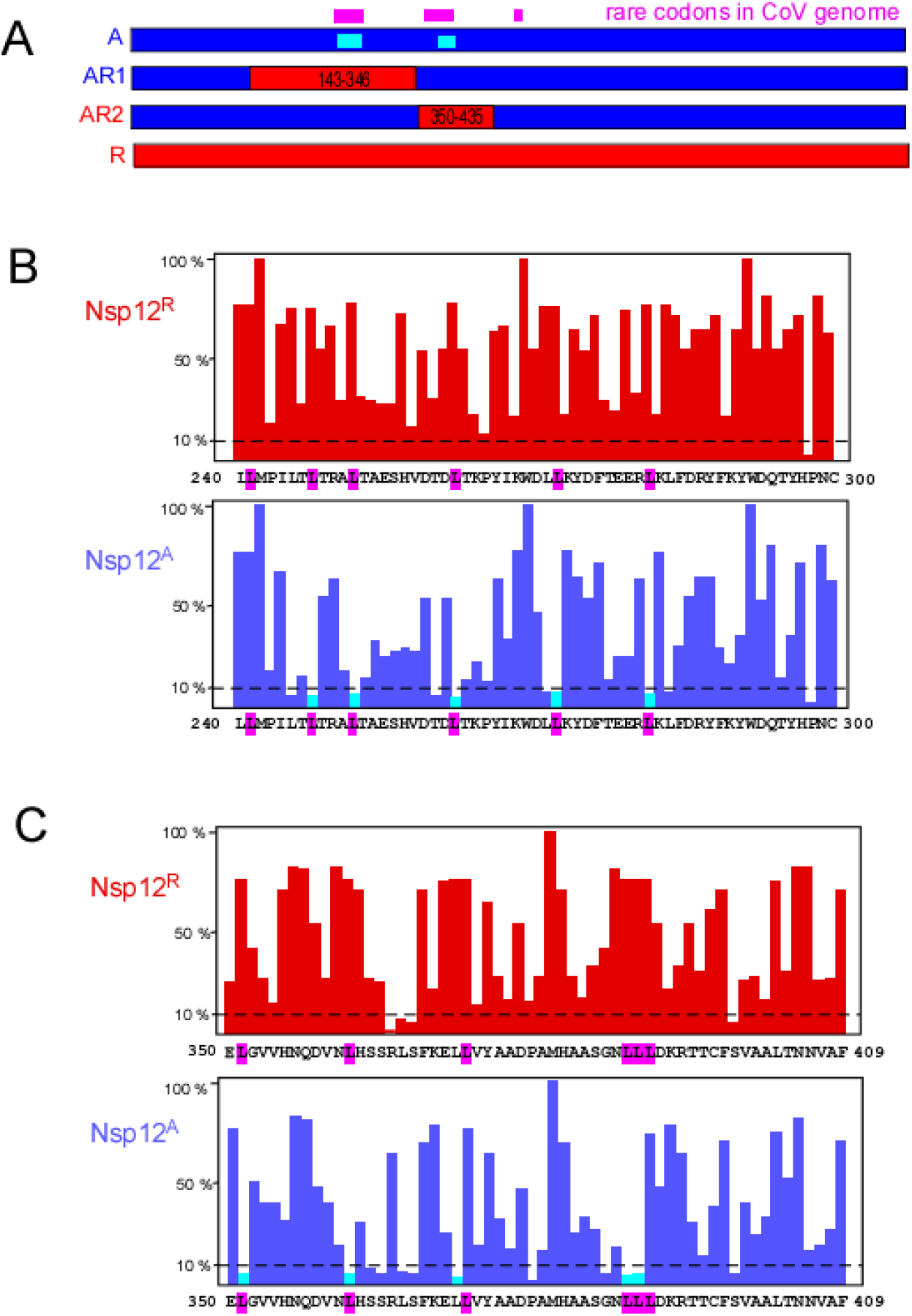
Rare codons in nsp12 CDS, as compared to codons present in highly expressed genes; this comparison is certainly appropriate for proteins expressed in *E. coli* from plasmids that carry a very strong T7 promoter and a canonical ribosome binding site. **A.** Linear maps of Nsp12 variants. Magenta bars on top show rare codons in the SARS CoV-2 genomic RNA; the cyan bars indicate regions in which rare codons at similar positions were also present in the Nsp12^A^, but not in the Nsp12^R^ coding sequence. **B** and **C.** The rare codon clusters in regions 1 (B) and 2 (C). The rare codons in the viral genome are highlighted on the sequence in magenta. The bar graph shows codon frequencies calculated using an on-line tool *E. coli* Codon Usage Analysis 2.0 developed by Morris Maduro (https://faculty.ucr.edu/~mmaduro/codonusage/usage.htm). A dashed line indicates the 10 % cut-off. Rare codons (below 10 %) present in the Nsp12A and viral CDS at identical or adjacent positions are shown in cyan.

**Figure S5.**
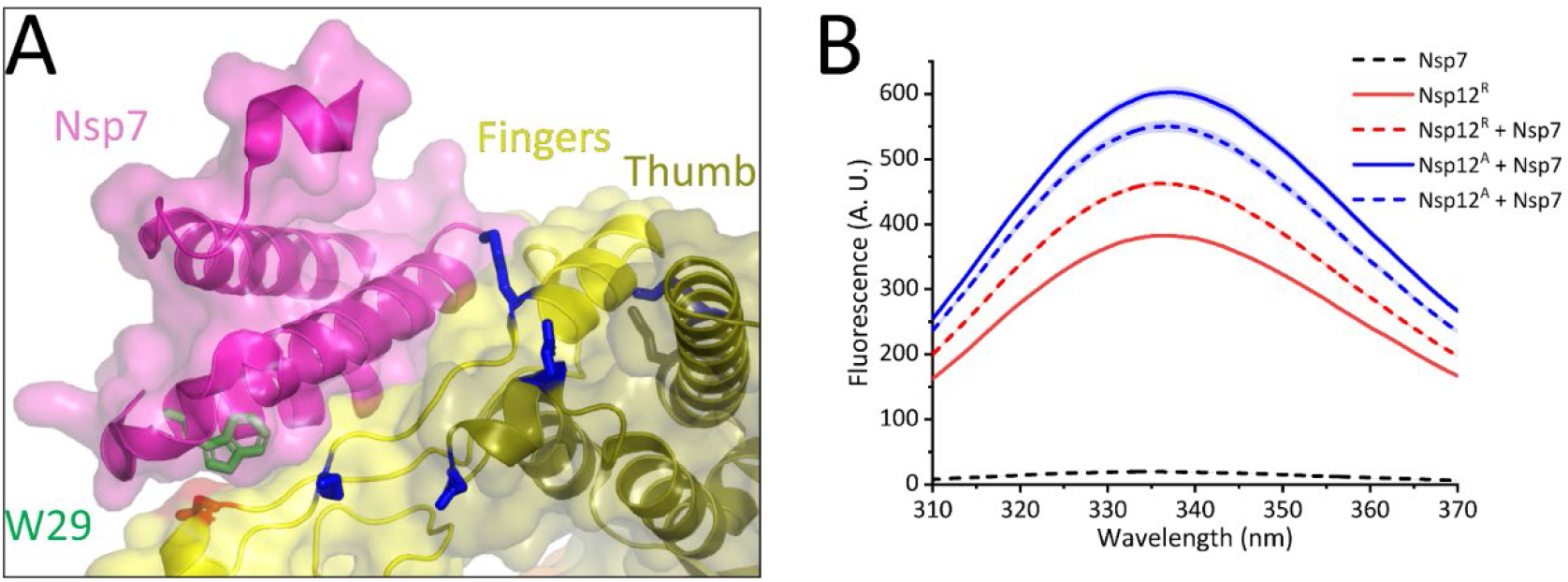
Nsp7/Nsp12 interactions. **A.** A zoom in at the Nsp7-12 interface, with positions of differential EDC modifications indicated as in **Figure 3B**. In Nsp7, a unique Trp29 residue is exposed but becomes buried at the interface with Nsp12. **B.** Nsp7/12 interactions assayed by Trp fluorescence. As expected, Nsp7 alone (with solvent-exposed Trp29) did not produce detectable emission. Upon addition of Nsp7 to Nsp12^A^, a modest decrease in intensity (relative to free Nsp12^A^) was observed, indicative of changes in both partners upon the complex formation, as docking of Nsp7 without concomitant changes in Nsp12 would be expected to increase fluorescence when Trp29 is buried. By contrast, an increase was observed when Nsp7 was added to Nsp12^R^. Most strikingly, although the emission spectra of the isolated Nsp12^R^ and Nsp12^A^ were very different, the spectra of Nsp7-Nsp12 complexes were tending to be similar. While we cannot identify specific Trp residues that become rearranged upon the complex formation, our results suggest that Nsp7 binds to both Nsp12 subunits.

